# Comprehensive profiling of antibiotic resistance, virulence genes, and mobile genetic elements in the gut microbiome of Tibetan antelopes

**DOI:** 10.1101/2025.07.24.666617

**Authors:** Jian Liu, Hong-Bo Ni, Ming-Yuan Yu, Si-Yuan Qin, Hany M. Elsheikha, Peng Peng, Li Guo, Lin-Hong Xie, Hong-Rui Liang, Cong-Cong Lei, Yu Xu, Yan Tang, Hai-Long Yu, Ya Qin, Jing Liu, Hong-Chao Sun, Xiao-Xuan Zhang, Bin Qiu

## Abstract

Tibetan antelopes, native to high-altitude plateau regions, play a vital role in the local ecosystem. However, their gut microbiota harbors diverse antibiotic resistance genes (ARGs) and virulence genes (VFGs), raising concerns about the potential spread of antibiotic resistance in wildlife. In this study, in addition to collecting 26,608 metagenome-assembled genomes (MAGs) from public databases, we performed gut metagenomic sequencing on 68 Tibetan antelopes and obtained 7,318 MAGs through binning. A total of 2,968 ARGs were identified, conferring resistance to 23 antibiotic classes, with elfamycin resistance being the most prevalent. Comparative analysis revealed 7 ARGs unique to Tibetan antelopes, suggesting species-specific adaptations. Additionally, during the analysis of shared antibiotic resistance profiles between Tibetan antelopes and humans, two bacterial strains were identified within the Tibetan antelope gut microbiota: *Enterococcus gallinarum* exhibiting resistance to extended-spectrum class A beta-lactamases and *Klebsiella grimontii* demonstrating vancomycin resistance. Mobile genetic elements played a key role in ARG dissemination. ARGs were significantly correlated with VFGs, particularly those linked to adherence and effector delivery systems. These findings underscore for the first time the potential ecological and health implications of ARG dissemination in Tibetan antelopes, highlighting the need for further surveillance to assess its impact on wildlife and environmental resistomes.

## Introduction

Antibiotics are among the most influential medical discoveries of the 20th century, having saved millions of lives by combating bacterial infections (1). However, the overuse and misuse of antibiotics have led to a steady rise in antibiotic resistance genes (ARGs), posing a serious global health threat. Since the early 21st century, the growing prevalence of antibiotic resistance has raised concerns about the diminishing effectiveness of current treatments, leaving humanity vulnerable to untreatable infections (2). The rapid spread of ARGs has become a critical public health crisis, with significant implications for both human and environmental health (3).

ARGs are naturally present in diverse ecosystems, where they serve as a defense mechanism against antibiotics produced by other microorganisms (4–6). The origins of resistance genes can be traced back millions—even billions—of years, yet their widespread dissemination has accelerated due to anthropogenic influences (6, 7). Through horizontal gene transfer, ARGs can spread across microbial communities, creating reservoirs of resistant bacteria (5). While antibiotic resistance has historically been linked to agricultural and healthcare settings, recent studies indicate that ARGs are now prevalent even in remote natural environments. This suggests that indirect human activities and environmental factors may contribute to their dissemination (8).

Beyond human and environmental influences, animals also play a crucial role in the spread of ARGs. Wildlife, particularly species in isolated or extreme habitats, may serve as unexpected reservoirs of antibiotic resistance, facilitating the movement of ARGs beyond human-dominated landscapes (9). Understanding how natural environments contribute to or mitigate the spread of resistance genes is essential for addressing the broader implications of antibiotic resistance.

The Qinghai-Tibet Plateau (QTP), often referred to as the "Roof of the World," is one of Earth’s most extreme and ecologically unique regions, averaging over 4,000 meters in altitude. This vast, high-altitude plateau spans western China, bordering India in the west and the Kunlun, Arjin, and Qilian Mountains in the northeast and northwest (10). The plateau’s harsh conditions—low oxygen levels, intense ultraviolet radiation, and extreme cold—have led to the evolution of highly specialized flora and fauna adapted to survive in this environment (11). Despite its relative isolation, the QTP is increasingly exposed to human activities such as livestock grazing, tourism, and antibiotic use in agriculture, which may introduce antibiotic residues and resistant bacteria into the ecosystem. These unique ecological conditions provide an exceptional opportunity to study the distribution and persistence of ARGs in a natural setting, as well as their potential transmission within and between wildlife species.

The Tibetan antelope (*Pantholops hodgsonii*), an iconic species endemic to the QTP, is of particular interest in this study due to its ecological significance and conservation status.

Listed in the Convention on International Trade in Endangered Species (CITES) since 1981, the Tibetan antelope has experienced population recovery in recent years and is now classified on the IUCN Red List of Endangered Species (12, 13). Adapted to the extreme conditions of the plateau, this species plays a key role in the local ecosystem. Given its exposure to diverse microbial communities, the Tibetan antelope may act as a reservoir or vector for ARGs, contributing to their distribution across the plateau. However, research on ARG dynamics in Tibetan antelopes remains limited, particularly in comparison to other plateau-dwelling species.

This study aims to investigate the role of Tibetan antelopes in the transmission and distribution of ARGs within the Qinghai-Tibet Plateau ecosystem. By analyzing bacterial diversity and ARG abundance in Tibetan antelopes and comparing these findings with data from five other plateau species, we seek to identify unique patterns of ARG transmission and assess the ecological risks associated with ARG-host relationships. Additionally, we examine the interactions between ARGs, mobile genetic elements (MGEs), and virulence genes (VFGs) to understand their potential role in the dissemination of antibiotic resistance in this extreme environment. Viruses, serving as important reservoirs of ARGs(14, 15), see their horizontal gene transfer (HGT) generally mediated by transduction via phages—viruses that infect bacteria(16). Therefore, predicting viral sequences and detecting ARGs were necessary to investigate the drug resistance profile in Tibetan antelopes.

The findings from this study will provide valuable insights into the ecological roles of wildlife in ARG spread and contribute to a broader understanding of antibiotic resistance in remote and extreme environments. This research not only establishes foundational knowledge about ARGs within Tibetan antelopes but also offers a new perspective on the mechanisms driving antibiotic resistance in high-altitude ecosystems. The results may have broader implications for global studies on ARG dissemination and wildlife-associated resistomes.

## Methods

### Tibetan antelopes sample collection

Freshly excreted feces of Tibetan antelopes were located using a telescope. Subsequently, fresh samples were carefully collected using sterile tweezers and transferred into labeled sterile centrifuge tubes. A total of 68 fecal samples were obtained from Tibetan antelopes across Tibet, Qinghai, and Xinjiang (Supplementary Table S1). All samples were stored at −80°C until further processing.

### Metagenomic sequencing and data analysis

Total DNA was extracted from the 68 fecal samples collected in this study using the OMEGA Mag-Bind Soil DNA Kit (M5635-02, Omega Bio-Tek, Norcross, GA, USA) following the manufacturer’s instructions. DNA concentration and purity were measured using a Qubit™ 4 Fluorometer and verified by 1% agarose gel electrophoresis. Samples meeting quality control standards were adjusted to a final concentration of 10 nM for library preparation. The sequencing libraries were constructed by Shanghai Personalbio Technology Co., Ltd. and sequenced on the Illumina HiSeq platform using paired-end (2 × 150) sequencing.

Fastp (v0.23.0) was used to filter for high-quality reads, which were then processed with Bowtie2 (v2.5.0) to remove host genomic (PHO1) DNA contamination. Contigs were assembled using MEGAHIT (v1.2.9) (17), and sequencing depth files were generated with BWA (v0.7.17-r1198) (18), SAMtools (v1.18) (19), and the script jgi_summarize_BAM_contig_depths. Binning was conducted with MetaBAT2 (v2.15) (20) using the parameters -m 2000 -s 200000 --seed 2023. The completeness and contamination of each bin were assessed using CheckM2 (v1.0.1) (21), retaining bins with completeness ≥50% and contamination ≤10%.

This study obtained 7,386 MAGs and 26,607 public Tibetan antelope MAGs were dereplication at 99% Average Nucleotide Identity (ANI) was performed using dRep (v3.4.3) (22) with the parameters -pa 0.9 -sa 0.99. Taxonomic classification of the dereplicated metagenome-assembled genomes (MAGs) (99% identity clusters) was performed using GTDB-Tk with default parameters (23), referencing the Genome Taxonomy Database (GTDB) to assign taxonomic ranks from domain to species. phylogenetic trees were constructed at the amino acid level using PhyloPhlAn (v.1.0).

### Public data collection and pre-processing of sequencing reads

In addition to the 68 fecal samples collected from Tibetan antelopes in this study and 255 Tibetan antelope samples and 26,607 MAGs were retrieved from CNP0001390 (24). Fecal metagenomic data from other herbivores were obtained from the China National GeneBank (CNGB) database. These inclused samples from *P. hodgsonii* and five other large herbivore species: yak (*Bos grunniens*, 28,125 MAGs), Tibetan wild ass (*Equus kiang*, 6,684 MAGs), Tibetan sheep (*Ovis aries*, 39,378 MAGs), Tibetan cattle (*Bos taurus*, 10,630 MAGs), and Tibetan horse (*Equus caballus*, 8,144 MAGs).

Furthermore, 60,664 high and medium quality MAGs of human (10.1038/s41586-019- 1058-x) were also obtained. Additionally, 32 fecal metagenomic datasets from Tibetan human individuals were obtained from NCBI, consisting of 17 samples from PRJEB53209 (25) and 15 from PRJNA543906 (26). All Tibetan humans fecal metagenomic datasets were processed using the same standardized data analysis pipeline as applied to the 68 fecal samples from Tibetan antelopes. For the Tibetan human samples, host-derived reads were removed by mapping against the GRCh38 reference genome using Bowtie2 (27) (v2.4.1) with default parameters.

### Identification and processing of viral sequences

To identify viruses potentially involved in the dissemination of AMR, established methods were used to screen contigs >5,000 bp from 2,149 MAGs carrying ARGs (^28, 29^). Initially, CheckV (v1.0.1) was used to assess the ratio of viral to host genes (^30^). Contigs containing more than 10 host genes or where host genes outnumbered viral genes by more than fivefold were excluded. Proviral fragments were also identified using CheckV. Next, multiple viral detection strategies were employed, including (1) viral gene enrichment as determined by CheckV, (2) identification by DeepVirFinder (v1.0.19) (^31^)with a score greater than 0.90 and p-value less than 0.01, and (3) viral classification by VIBRANT (v1.2.1) (^32^)using default parameters. Contigs meeting any of these criteria were retained as putative viral sequences.

To remove potential bacterial contamination, BUSCOs (^33^) were used in combination with hmmsearch to detect bacterial single-copy orthologs within the viral candidates. The BUSCO ratio (calculated as the number of BUSCOs divided by total gene count) was computed, and sequences with a ratio of 5% or higher were excluded. The remaining sequences underwent quality assessment with CheckV, and only viral genomes with medium or higher completeness were retained for downstream analyses.

### Functional annotation

Gene prediction for all MAGs obtained in this study was performed using Prodigal (10.1186/1471-2105-11-119) (v2.6.1). ARGs were identified by aligning protein sequences against the Comprehensive Antibiotic Resistance Database (CARD v3.2.7) (34) using DIAMOND (v2.1.8.162) (23), with a threshold of >80% sequence identity and >80% query coverage (e-value = 1e-5) (35). Multidrug resistance genes were defined as those conferring resistance to at least two antibiotic classes, while multitype mechanism genes conferred resistance via at least two distinct mechanisms. MGEs were detected by aligning gene sequences to the MGE Database (36) using BLASTN (v2.13.0) (37) with the parameters - evalue 1e-5 -perc_identity 80 -qcov_hsp_perc 80. VFGs were identified by aligning sequences to the Virulence Factor Database (VFDB) (38) using DIAMOND (v2.1.8.162), with a sequence identity threshold of >80% and query coverage >80%.

To generate taxonomic profiles for Tibetan antelope and human-derived ARGs, clean reads from each sample were mapped to reference ARG sequences using Bowtie2 (v2.4.1) with the parameters --end-to-end --fast --no-unal. Total mapped reads across all samples were normalized to the same sequencing depth. The read counts were normalized to transcripts per kilobase million (TPM).

### Statistical analyses and visualization

The resulting phylogenetic trees were annotated and visualized using iTOL (39). Procrustes association analysis was conducted on the profiles of ARGs, MGEs, and VFGs, utilizing the ’procrustes’ function from the ’vegan’ package. Taxonomic assignments of genomes containing ARGs were identified, and Sankey plots were generated using the ‘ggsankey’ package (v0.0.9). The Richness index was calculated from the relative abundance of functional genes. The correlation between ARGs, VFGs, and MGEs was assessed using the cor.test function in R, with Spearman’s rank correlation method. Gene arrow maps were constructed using the ‘gggenes’ (v0.4.1) package. GCView services (https://proksee.ca/) was used to visualize genome and to mark ARGs, MGEs, and VFGs. Network visualization of these correlations was conducted using the igraph v2.14.0 package. All other visualizations were produced with the ggplot2 package version 3.3.6. Statistical analyses were performed using R version 4.4.1.

## Results

### Comprehensive gut genome of the Tibetan antelope

A total of 33,925 MAGs from Tibetan antelope gut microbiota were obtained from public databases (see Methods). After quality control using CheckM2, 28,752 medium- to high-quality MAGs (completeness ≥ 50% and contamination ≤ 10%) were retained. These MAGs were clustered at 99% ANI, resulting in 13,600 strain-level genomes (Fig. 1). The average completeness of these genomes was 82.50%, with a mean contamination of 1.95% (Supplementary Fig.1B). Genome sizes ranged from 0.38 to 6.50 Mbp (average 1.89 Mbp/genome), while GC content varied between 23.93% and 73.57% (average 45.82%) (Supplementary Fig.1C).

**Fig. 1.**
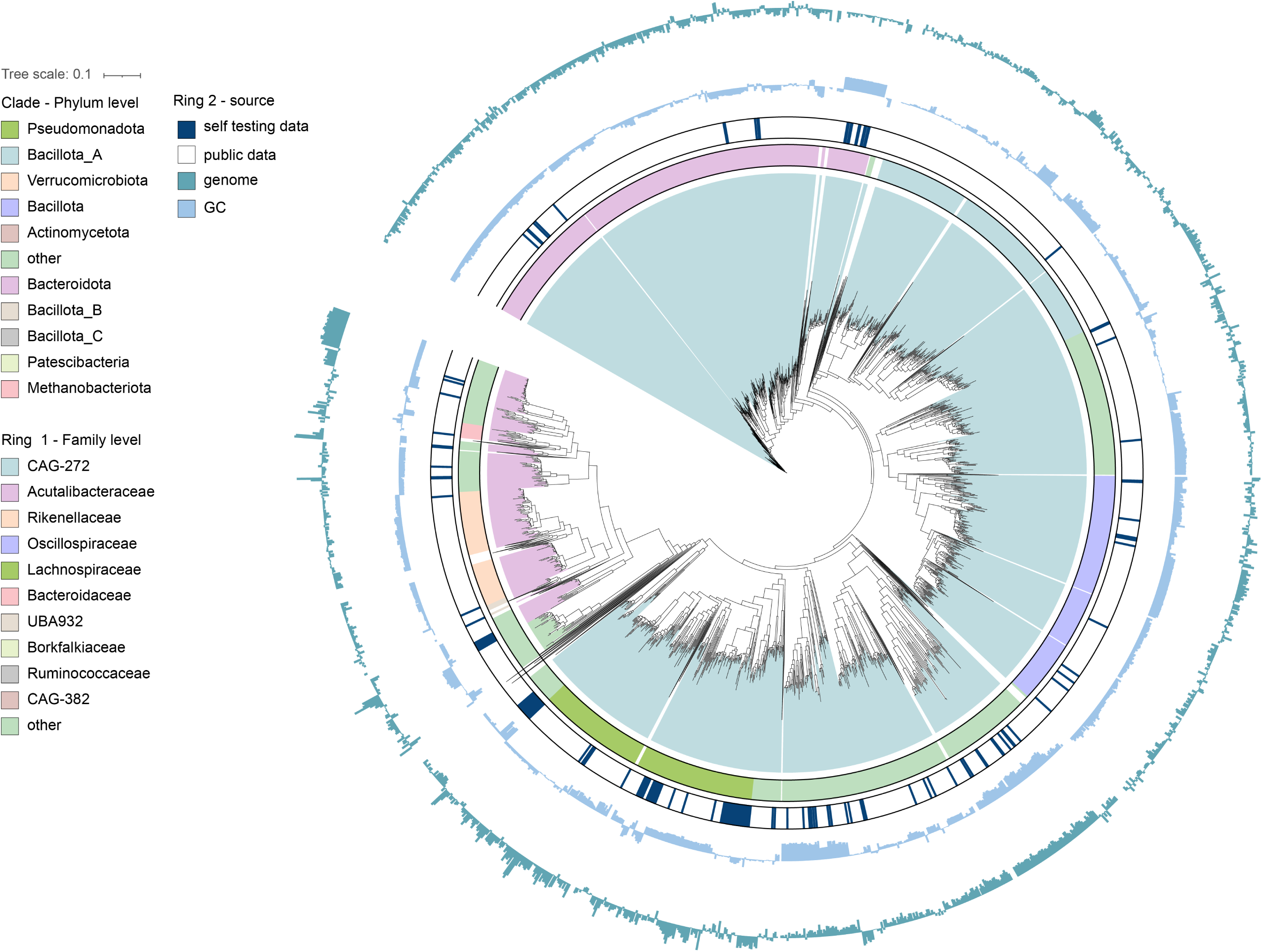
Metagenome-assembled genomes (MAGs) of Tibetan antelope. The phylogenetic relationship among the 13,600 bacterial genomes. The color coding of each clade corresponds to the phylum-level classification of the genomes. The first outer ring denotes family level classification. The second outer ring denotes the genome source, distinguishing between self data and public data. The third and fourth rings are bar charts representing the GC content and the genome size of each genome, respectively.

Taxonomic classification using GTDB-Tk assigned these genomes to 2 domains, 21 phyla, 28 classes, 76 orders, 164 families, and 674 genera. At the domain level, 14,234 genomes were classified as bacteria, while 335 genomes belonged to archaea (Supplementary Table S2). The dominant phylum was *Bacteroidota_A* (7,963 genomes, 58.55%), followed by *Bacteroidota* (3,488 genomes, 25.65%). At the genus level, *Alistipes* had the highest representation (1,208 genomes, 8.88%), followed by *Faecousia*, *Cryptobacteroides*, and *Scatosoma* (Supplementary Fig.1D). Notably, 343 genomes (2.52%) remained unclassified at the genus level, indicating the presence of potentially novel microbial taxa in the Tibetan antelope gut microbiome.

### Composition of ARGs in Tibetan antelope

We compared 13,600 genomes to the CARD and identified 2,968 ARGs. These genes confer resistance to 23 different antibiotics (Supplementary Table S3). The ARGs were classified into five resistance mechanisms, excluding multi-drug resistance. Antibiotic target alteration was the most prevalent mechanism, present in 2,478 genomes (83.49% of all resistance mechanisms). The least common mechanism was reduced permeability to antibiotics, found in only 12 genomes (0.40%). Multi-drug resistance ranked second, with 212 genomes (7.14%) exhibiting this mechanism (Supplementary Table S3).

We categorized the types of antibiotics these resistance genes target. The most prevalent resistance was against elfamycin antibiotics, comprising 56.44% of the total, followed by multi-type drug resistance (14.25%), glycopeptide antibiotics (8.83%), fusidane antibiotics (7.28%), and peptide antibiotics (4.08%). The least common resistance was to phosphonic acid antibiotics, found in 31 genomes (1.04%) (Fig. 2A).

**Fig. 2.**
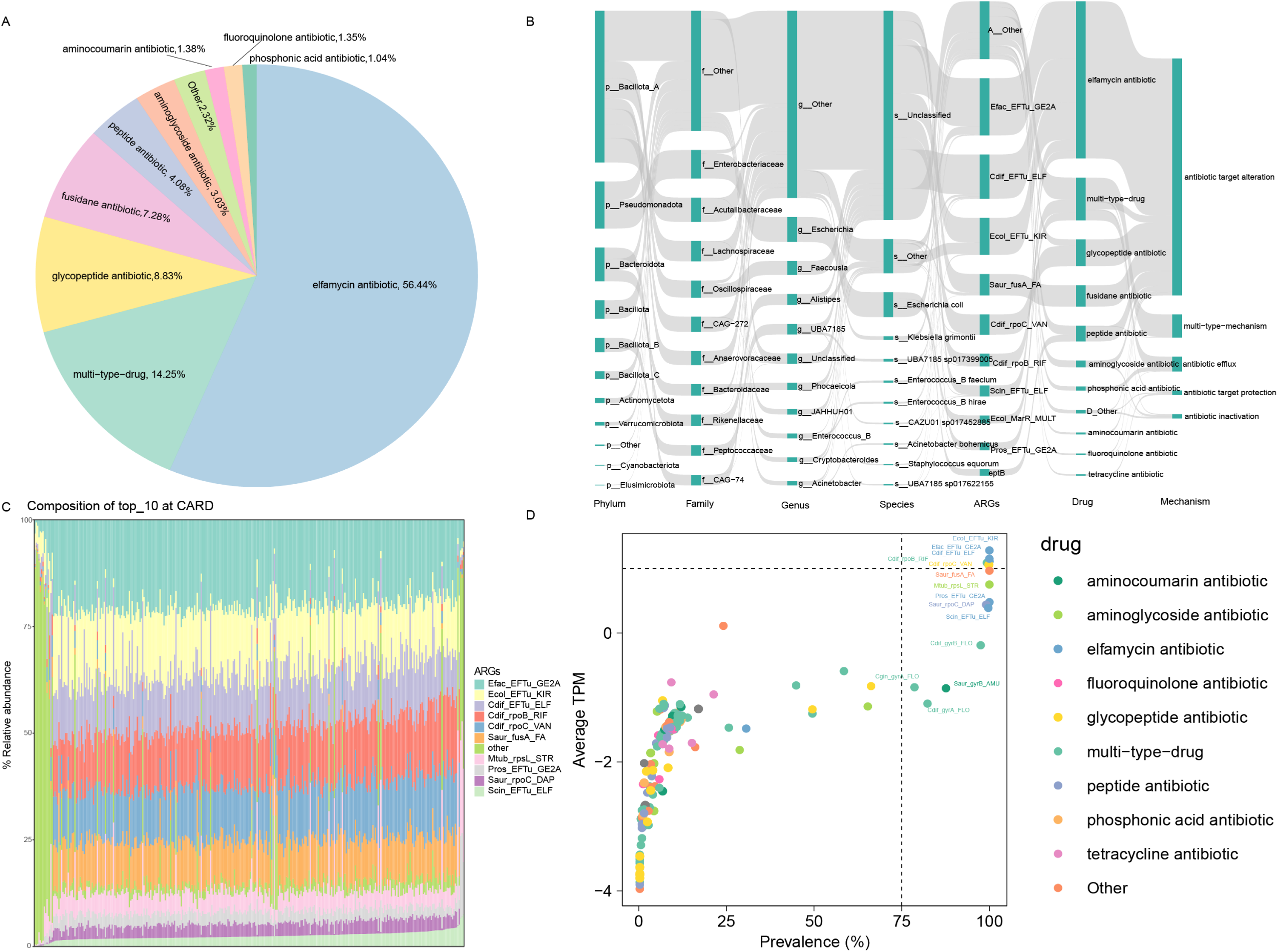
Composition of ARGs in Tibetan antelope. (A) Proportion of resistance to different drug classes. (B) Host classification of ARGs, including their resistance mechanisms. (C) Composition of the top 10 ARGs. (D) Prevalence and abundance distribution of all ARGs in the 323 tested samples. The abundance of each ARG is represented as the mean value across all samples. Dots of different colors represent distribution patterns based on abundance levels and frequencies.

Host prediction analysis at the phylum level revealed that *Bacillota_A* contained the highest number of ARGs, followed by *Pseudomonadota*, *Bacteroidota*, *Bacillota*, and *Bacillota_B*. At the species level, a significant proportion of ARGs could not be clearly classified. Among those that were classified (excluding unresolvable species), *Escherichia coli* harbored the largest number of ARGs, particularly resistant to peptide antibiotics. In addition to this, *E. coli* exhibited resistance to multiple other antibiotics as well (Fig. 2B).

To further investigate the composition of antibiotic resistance genes (ARGs), we examined the distribution of the top 10 ARGs in each Tibetan antelope sample (Fig. 2C). In over 75% of the samples, the proportion of *Efac_EFTu_GE2A* exceeded 12.5%. This was followed by *Efac_EFTu_KIR* and *Cdif_EF_ELF*, with the combined proportion of these three ARGs surpassing 50% in nearly half of the samples.

The top 10 ARGs confer resistance to five types of antibiotics: elfamycin, peptide, rifamycin, glycopeptide, and aminoglycoside antibiotics. The primary resistance was observed towards elfamycin antibiotics, with genes such as *Ecol_EFTu_KIR*, *Cdif_EFTu_ELF*, *Pros_EFTu_GE2A*, and *Scin_EFTu_ELF* contributing to this resistance. Additionally, *Cdif_rpoB_RIF* provides resistance to both peptide antibiotics and rifamycin antibiotics. Except for *Cdif_rpoB_RIF*, all other ARGs primarily exhibit resistance through antibiotic target alteration. *Cdif_rpoB_RIF*, however, involves both antibiotic target alteration and a secondary mechanism of antibiotic target replacement. The prevalence of these ARGs was detected in over 75% of the samples (Fig. 2D).

### ARGs diversity across Tibetan antelope, Human, and other Tibetan mammals

The methods for processing the MAGs of other plateau animals, including Tibetan wild asses, yaks, Tibetan wild horses, and Tibetan wild cattle were consistent with those used for Tibetan antelope MAGs. Among the plateau animals, Tibetan sheep had the highest number of MAGs, with 39,378 MAGs collected, and the largest number of ARGs mapped to them (5,961 ARGs). This was followed by Tibetan yak (4,935 ARGs) and Tibetan wild cattle (2,009 AGRs) (Supplementary Table S4-S8). Further analysis of ARGs in these plateau-dwelling animals and human populations revealed that, excluding those categorized as ’others’, the top three most prevalent ARGs were primarily associated with multi-type-drug resistance, peptide antibiotics, and phosphonic acid antibiotics (Fig. 3A-B).

**Fig. 3.**
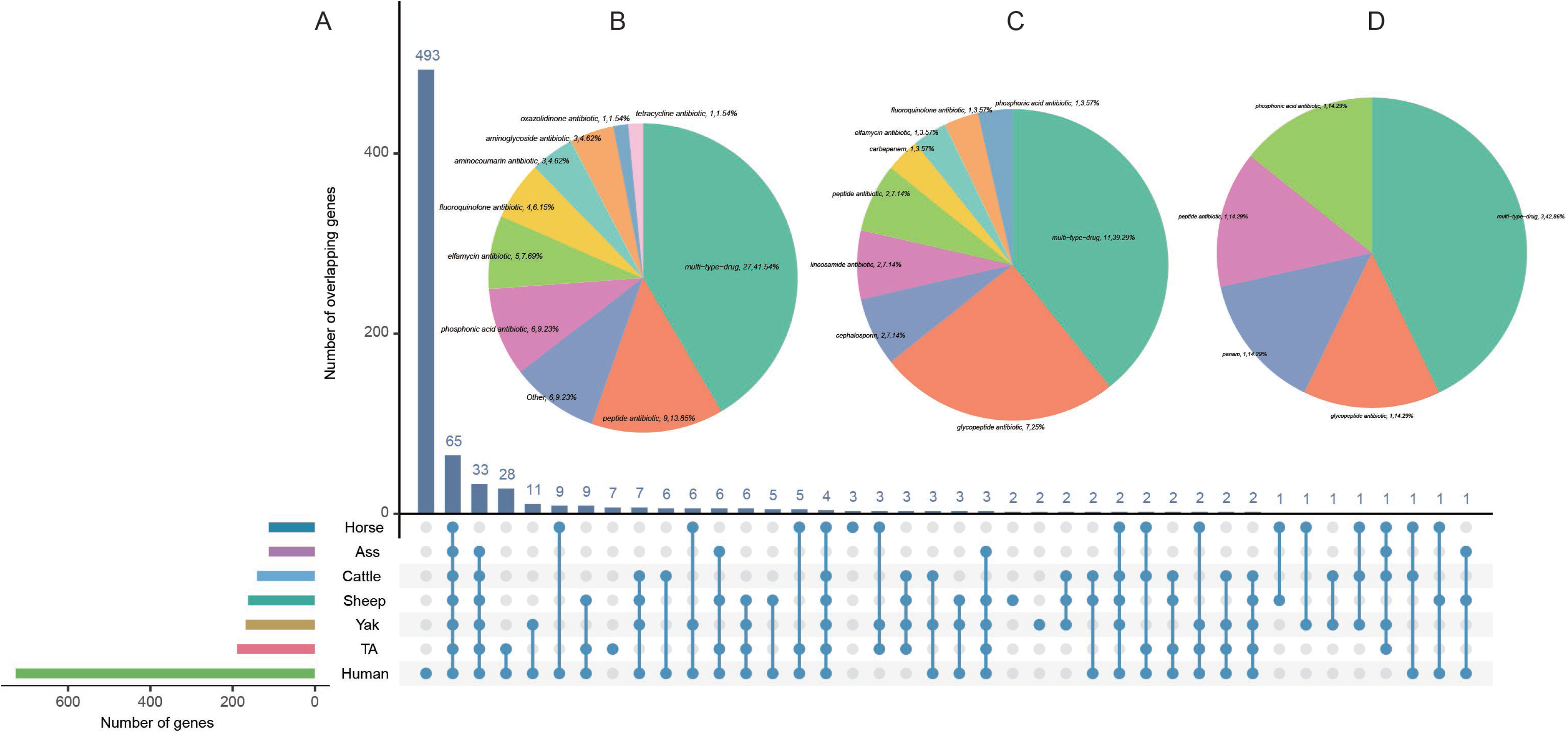
MAGs and ARGs analysis of other Tibetan mammals and human. (A) Comparison of ARGs between Tibetan antelopes and other species, with the vertical axis representing common ARGs across species. (B) Shared ARGs between plateau-dwelling animals and human populations. (C) Shared ARGs between Tibetan antelope and human populations. (D) ARG types exclusively present in the gut microbiota of Tibetan antelopes.

Comparative analysis of shared ARGs between Tibetan antelopes and human populations identified 28 distinct ARG types. Among these, 11 ARGs conferred resistance to multiple drug classes, while 7 ARGs specifically targeted glycopeptide antibiotics (Fig. 3C). Notably, we identified 7 ARGs exclusively present in Tibetan antelopes, with no counterparts detected in other plateau-dwelling species or Tibetan populations. Particularly significant were three multidrug-resistant ARGs in Tibetan antelopes demonstrating simultaneous resistance to carbapenems, cephalosporins, and penams. Subsequently, BLAST analysis of the shared antibiotic resistance genes (ARGs) between Tibetan antelopes and humans via NCBI identified a vancomycin-resistant Enterococcus gallinarum strain and an extended-spectrum class A beta-lactamase (ESBL)-producing Klebsiella grimontii strain (Supplementary Fig. 2A, Supplementary Table S10).

### MGEs related to ARGs in the gut of Tibetan antelope

MGEs play a critical role in the horizontal transfer of ARGs, both within bacterial populations and across different species. Understanding their distribution patterns and their relationship with ARGs is essential for comprehending the mechanisms of antibiotic resistance spread. In this study, a total of 132 MGEs were identified from 13,600 MAGs by aligning protein sequences with the MGE database. These MGEs were classified into five categories: Transposon, Insertion_element, and others (Supplementary Table S11).

Transposons, characterized by transposase genes, were the most abundant MGE type in the Tibetan antelope gut microbiome, comprising 46.52% of the total MGE abundance. This was followed by Insertion_element (43.32%), integrons (7.03%), Plasmids (2.89%), and unknown elements (0.25%) (Fig. 4A).

**Fig. 4.**
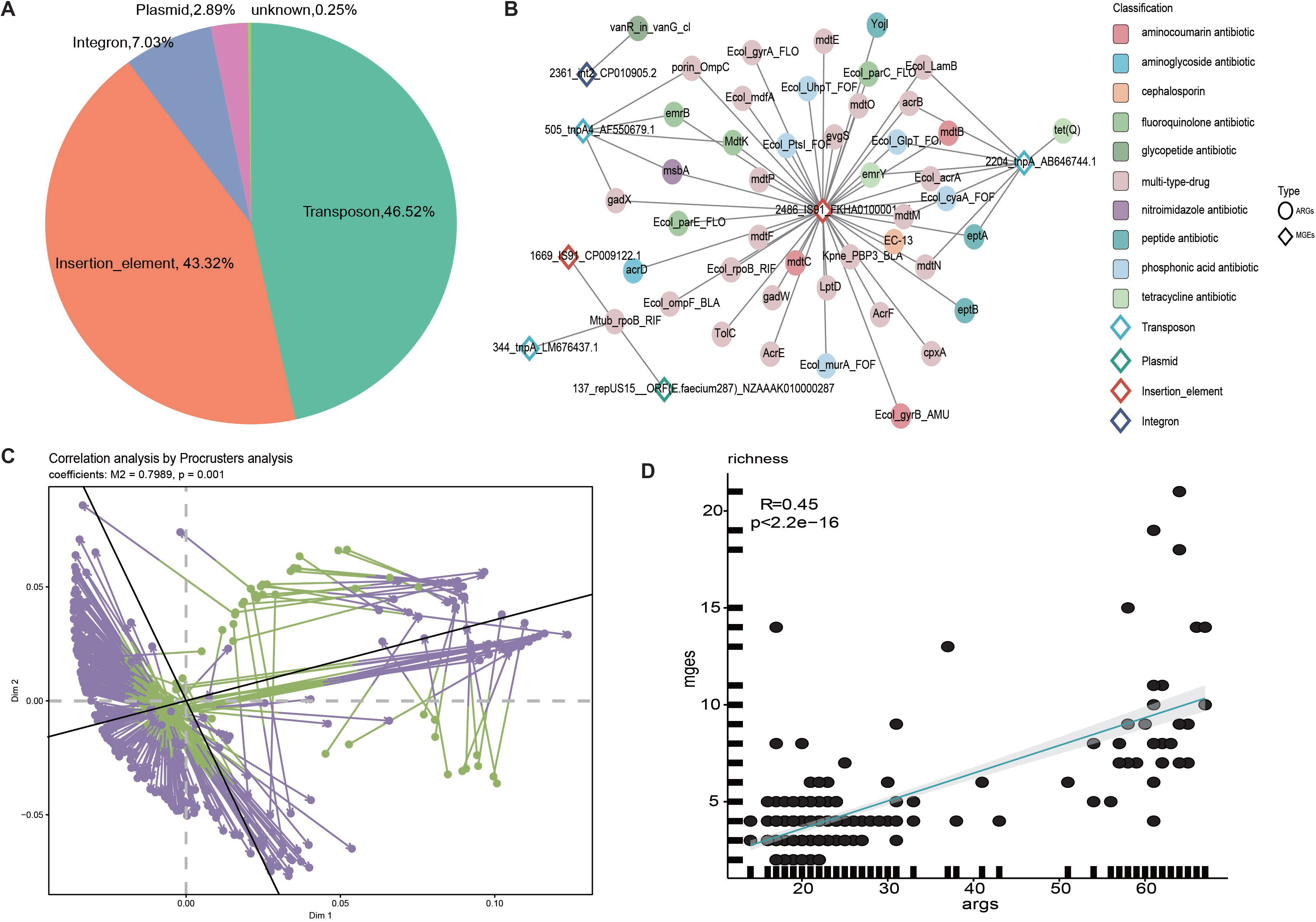
Mobile genetic elements (MGEs) related to ARGs. (A) Abundance of different types of MGEs across samples. (B) Correlation network between MGEs and ARGs. ARGs are represented by circles, and MGEs by rhombuses. (C) Procrustes association analysis: Correlations between ARGs and MGEs in multidimensional space. (D) Spearman’s correlation analysis of the richness index between MGEs and ARGs.

We then conducted a correlation analysis between the MGEs and ARGs, focusing on relationships with a corr > 0.6 and a *p*-value < 0.05 (Fig. 4B). The analysis showed that Insertion_elements were associated with a higher number of ARGs compared to other MGE types. Notably, the majority of ARGs linked to these MGEs were related to aminocoumarin antibiotics. Further, procrustes analysis revealed a strong association between the cecal mobilome and antibiotic resistance (PROTEST, M² = 0.7989, *p* = 0.001) (Fig. 4C). This was supported by a significant positive correlation between the MGE and ARG profiles, as measured by the Richness index (Spearman’s correlation: R = 0.45, *p* < 2.2^e-16^) (Fig. 4D).

Additionally, ARGs located within 5 kilobases (kb) of an MGE were classified as potentially mobile. Only one Escherichia coli strain (X23.bin.1) was identified carrying two ARG-MGE configurations: Ecol_emrE (conferring resistance to macrolide antibiotics) paired with ISSfl3, and AcrF (associated with multi-type-drug resistance) linked to IS91. Notably, the AcrF gene demonstrated resistance to cephalosporins, cephamycins, fluoroquinolone antibiotics, and penams (Supplementary Fig.2B). Therefore, these two resistance genes are considered potentially mobilizable.

Additionally, some MAGs contained phage sequences. Viral sequence prediction was performed on the MAGs carrying ARGs, identifying a total of 7,802 viral sequences.

Following clustering and annotation, the majority of these sequences were found to belong to the *Siphoviridae* family. These clustered viral sequences were then aligned against the CARD database, revealing two viral sequences (taxonomic family is unknown) harboring ARGs.

These ARGs conferred resistance to fusidane antibiotics and lincosamide antibiotics, respectively (Supplementary S3).

### Composition of VFGs and their relationship with ARGs in Tibetan antelope

To further analyze the composition of VFGs in the gut metagenome of Tibetan antelopes, we identified 4,729 VFGs by comparing the metagenomes with the VFDB. Among these, the most abundant virulence factor gene was *tufA*, which accounted for 48.18% of the VFGs (Supplementary Table S14). This gene was present in nearly all Tibetan antelope samples, with its relative abundance exceeding 25% in most cases (Fig. 5A). The second most prevalent VFG was *cps4L*, accounting for 19.86%, followed by *groEL* at 12.06%, *cps4J* at 9.85%, and *sigA/rpoV* at 1.43%. Based on functional classifications, the two virulence factor genes *tufA* and *groEL* are primarily associated with adherence functions, while *cps4L* and *cps4J* are linked to immune modulation. sigA/rpoV, on the other hand, is involved in regulatory functions (Supplementary Table S14).

**Fig. 5.**
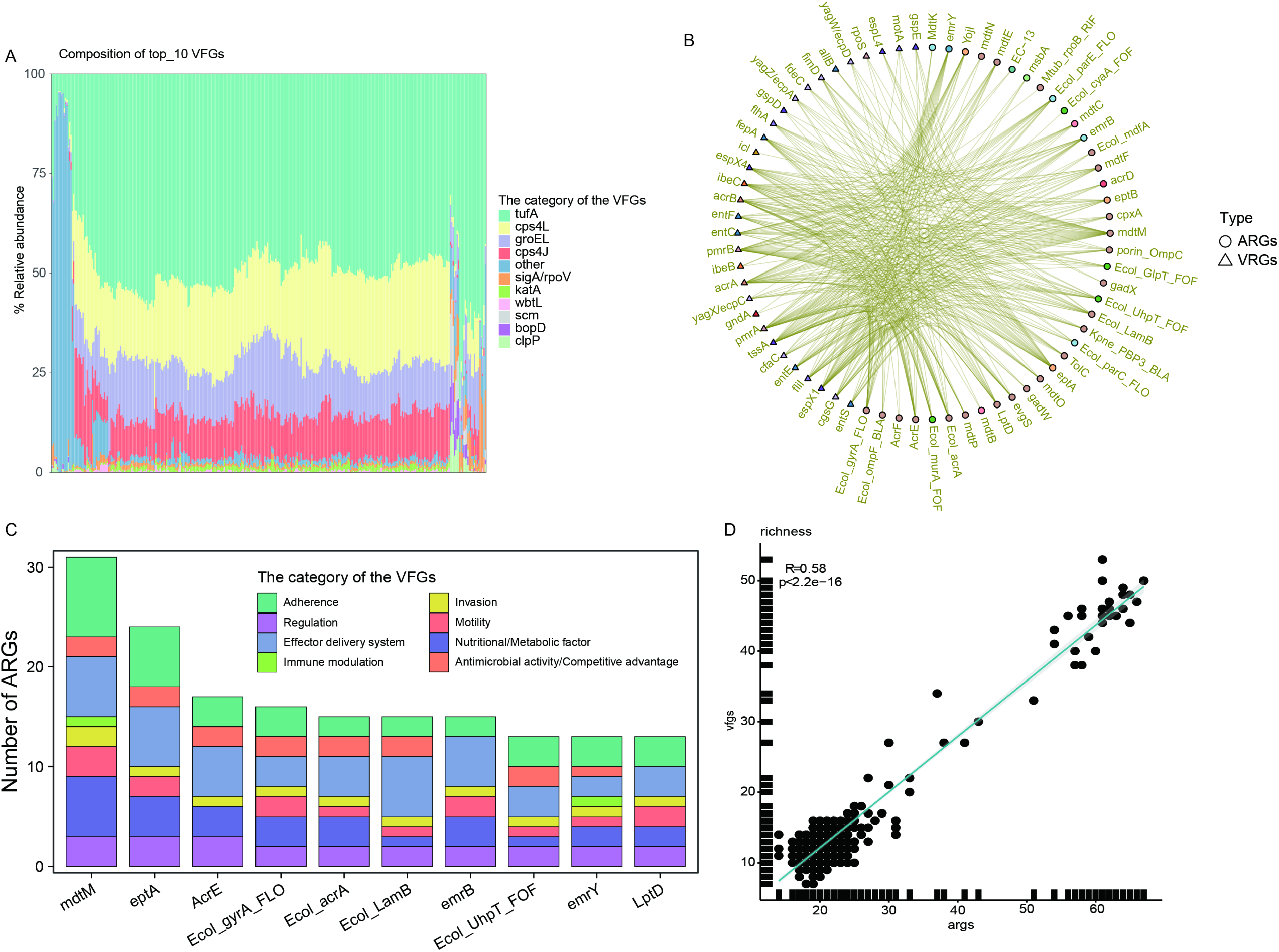
Composition of VFGs and correlation between ARGs and VFGs. (A) Abundance of VFs in Tibetan antelope samples. (B) Correlation between ARGs and VFGs, with corr > 0.8 and *p*-value < 0.05. (C) Number of correlations for the top 10 ARGs, with bars representing the number of each category of VFGs associated with the ARGs. (D) Spearman’s correlation analysis between the richness index of VFGs and ARGs.

To examine the relationship between ARGs and VFGs, we performed a correlation analysis using the Spearman’s method. We focused on relationships with a correlation coefficient (corr) > 0.8 and a *p*-value < 0.05 (Fig. 5B). We found that all the top 10 ARGs showed significant associations with VFGs related to adherence functions, with the highest number of such associations (Fig. 5C). The second most common association was with virulence genes related to the effector delivery system. Additionally, several of these ARGs confer resistance to multiple drugs, such as *mdtM*, *Ecol_acrA*, *Ecol_gyrA_FLO*, *Ecol_LamB*, and LptD (Supplementary Table S4). Our study revealed a significant positive correlation between the profiles of VFGs and ARGs based on the Richness index (Spearman’s correlation: R = 0.58, *p* < 2.2^e-16^) (Fig. 5D). Applying the aforementioned analytical approach to investigate associations between VFGs and MGEs, we identified a total of five distinct MGE-VFG configurations across two *Escherichia coli* strains (Supplementary Fig. 2C-E).

Notably, both strains harbored over 65 antimicrobial resistance genes (ARGs) each, a characteristic that may have significant public health implications.

## Discussion

Despite significant progress in understanding AMR, knowledge of its impact on wild animals, especially those living in plateau regions, remains limited. In this study, we explored the distribution patterns of ARGs within plateau-region wildlife, with a focus on the Tibetan antelopes. Additionally, we conducted thorough correlation analyses between the detected ARGs, and VFs and MGEs to better understand potential co-occurrence relationships.

Emerging evidence suggests that ARGs carried by wildlife may spread through horizontal gene transfer, which could contribute to the development of multidrug-resistant pathogens (40–42). After rigorous quality control and clustering procedures, we obtained 13,600 MAGs from the Tibetan antelope gut microbiota. Taxonomic annotation using GTDB-Tk revealed that many these MAGs (*n* = 13,600) remained unclassified at the species level, underscoring significant gaps in our current understanding of the microbial composition within the Tibetan antelope intestinal ecosystem.

Our analysis identified 2,968 ARGs across 13,600 MAGs, conferring resistance to 23 antimicrobial classes. Among these classes, aminoglycosides, tetracyclines, and fluoroquinolones are clinically prevalent, while elfamycin, aminocoumarins, and peptide antibiotics are less commonly encountered in clinical practice (43, 44). Furthermore, an abundance analysis revealed a high prevalence of resistance genes against elfamycin and glycopeptide antibiotics in the gut microbiota of Tibetan antelopes. Interestingly, except for those conferring resistance to aminoglycosides, the most prevalent ARGs primarily confer resistance to antibiotics rarely used in clinical settings.

Elfamycins, a class of naturally occurring antibiotics, primarily target elongation factor EF-Tu, a crucial GTPase in bacterial protein synthesis (45). EF-Tu is responsible for accurately delivering aminoacyl-tRNA to the ribosome’s A-site, ensuring the fidelity and efficiency of translation (46). Additionally, we found that the predominant resistance mechanism of these highly prevalent ARGs is antibiotic target alteration, a common resistance mechanism. This alteration typically arises from spontaneous mutations in bacterial chromosomal genes, compounded by selection pressure from the presence of antibiotics (47).

The high prevalence of ARGs against elfamycin and glycopeptide antibiotics in the Tibetan antelope gut microbiota underscores a potential natural reservoir of resistance genes that could be transferred to other microbial communities. This finding suggests that the microbial ecosystems of wild animals might serve as a source of resistance genes, particularly those conferring resistance to antibiotics less commonly used in clinical practice. While this may not pose an immediate clinical threat, it highlights the need for continuous surveillance of antibiotic resistance in wildlife populations, as these environments could harbor diverse and novel resistance mechanisms (48, 49). Furthermore, these ARGs may have been acquired through natural selection pressures present in wild environments (50). However, the detection of clinically relevant ARGs and antibiotic-resistant bacteria (ARB) in wildlife species without antibiotic exposure should be regarded as an indicator of environmental resistance pollution(51, 52).

We conducted a correlation analysis between ARGs, VFGs, and MGEs. Regarding the correlation between ARGs and VFGs, we found that the top ten ARGs in terms of correlation were mostly cross-resistant, primarily associated with virulence factors related to adherence and effector delivery systems. Additionally, we identified 2 MAGs with complex resistance genes and virulence factors. Upon examining their species annotations, we found they were identified as *E. coli*. These findings suggest that these species may harbor a significant combination of both antibiotic resistance and virulence traits, potentially enhancing their pathogenicity and potentially complicating treatment options (53). The emergence of multidrug resistance in *Escherichia coli* has become a growing global concern, with its prevalence increasingly observed not only in human medicine but also across veterinary practice(54).

The presence of cross-resistant ARGs, especially when coupled with virulence factors such as adherence and effector delivery systems, poses a significant threat to human health. These factors not only allow bacteria to resist commonly used antibiotics, making infections harder to treat, but also enhance their ability to adhere to host tissues and invade cells. This increases the severity and persistence of infections. The combination of antibiotic resistance and increased virulence may lead to more severe, difficult-to-treat infections and contribute to the spread of multidrug-resistant pathogens in clinical settings, presenting a major public health concern (55, 56).

The correlation analysis between ARGs and MGEs revealed that the ARGs conferring resistance to aminoglycoside antibiotics were predominantly associated with insertion elements, a type of MGE. Insertion sequences (IS) are the simplest mobile genetic elements within cells and can appear in high numbers in prokaryotic genomes (56). They play a crucial role in evolution by promoting genomic plasticity, facilitating genetic rearrangements, and enabling the horizontal transfer of genes, including those conferring antibiotic resistance (57, 58). This suggests that IS elements may play a key role in the dissemination of aminoglycoside resistance genes, potentially influencing the spread of antibiotic resistance within microbial communities.

Viruses represent one of the most diverse life forms on Earth(15) and profoundly influence biogeochemical cycles along with the ecology and evolution of microbial communities(59, 60). For instance, recent research demonstrates that three ARGs –*bla*TEM, *sul1* and *tetW* – carried by phage particles can amplify and disseminate within *Escherichia coli* cells, further confirming the dominance of ARG-bearing phage particles in food retail-sourced viromes(61). However, this study identified limited ARGs through alignment with predicted viral sequences, and these two sequences remain taxonomically unclassified. These ARGs conferred resistance to fusidane antibiotics and lincosamide antibiotics. Fusidane-type antibiotics, represented by helvolic acid, fusidic acid and cephalosporin P1, are fungi-derived antimicrobials with little cross-resistance to commonly used antibiotics(62). Compounded by the current scarcity of knowledge regarding viral-associated ARGs – particularly the contribution mechanisms of viruses to ARG horizontal gene transfer – these limitations pose significant challenges to our further investigation.

We compared the ARGs from Tibetan antelopes with those from other plateau animals and human population. We identified 65 shared resistance genes, the majority of which conferred resistance to muti-type-drug. Additionally, we found 7 unique ARG types in Tibetan antelopes that were not present in other species. Most of these ARGs exhibited cross-resistance, followed by resistance to glycopeptide antibiotics. This suggests that Tibetan antelopes may harbor a distinct resistome, potentially shaped by their unique habitat and microbial ecology(63). The high prevalence of elfamycin resistance genes across multiple species indicates a common selective pressure(64), while the distinct resistome of Tibetan antelopes suggests potential host-specific adaptations.

Our study has some limitations. For example, the data collection from different plateau animals and Tibetan residents was not conducted in the same region, which limits our ability to analyze the specific transmission of resistance between them. Additionally, the limited collection of gut metagenomic data from Tibetan residents may lead to deviations from the true resistance landscape in the plateau region. More comprehensive and regionally consistent data will be essential for providing a clearer understanding of the dynamics of antibiotic resistance in both Tibetan residents and plateau animals.

## Conclusion

We conducted a comprehensive analysis of antibiotic resistance genes (ARGs) in Tibetan antelopes and untangled key findings. A high prevalence of resistance to elfamycin, glycopeptides, and aminoglycoside antibiotics was observed in the gut microbiota of Tibetan antelopes. These animals exhibited a unique combination of resistance genes, including 7 distinct ARGs, most of which are associated with cross-resistance. The top ten most abundant ARGs were linked to virulence factors, suggesting enhanced pathogenicity. Mobile genetic elements (MGEs), especially insertion sequences, played a pivotal role in ARG transfer, showing a strong correlation between MGEs and ARG distribution. This association between ARGs and virulence factors indicates that Tibetan antelopes may face a higher risk of infection. Finally, we identified two viral sequences carrying ARGs that confer resistance to fusidane antibiotics and lincosamide antibiotics, respectively. Our findings highlight the complex interplay between resistance, virulence, and ecological factors. They underscore the need for further investigation into antibiotic resistance in wildlife and its implications for conservation and public health in plateau regions.

## Data availability

The sequencing reads from each sequencing library have been deposited in CNGB and NCBI under the accession numbers: CNP0001390, PRJNA1257558, PRJEB53209 and PRJNA543906. All supplementary figures and tables are provided as additional files.

## Author Contributions

Jian Liu: Writing -original draft, Formal analysis, Software, Visualization. Hong-Bo Ni: Project administration, Supervision, Writing-review and editing. Ming-Yuan Yu: Conceptualization, Resources, Writing-review and editing. Si-Yuan Qin: Resources, Supervision, Visualization, Writing-review and editing. Hany M. Elsheikha: Conceptualization, Validation, Writing-review and editing. Peng Peng: Funding acquisition, Resources, Writing-review and editing. Li Guo: Conceptualization, Writing-review and editing. Lin-Hong Xie: Resources, Writing-review and editing. Hong-Rui Liang: Resources, Writing-review and editing. Cong-Cong Lei: Resources, Writing-review and editing. Yu Xu: Resources, Writing-review and editing. Yan Tang: Validation, Supervision, Software, Writing-review and editing. Hai-Long Yu: Formal analysis, Writing-review and editing. Ya Qin: Resources, Writing-review and editing. Jing Liu: Conceptualization, Validation, Writing-review and editing. Hong-Chao Sun: Conceptualization, Formal analysis, Writing-review and editing. Xiao-Xuan Zhang: Conceptualization, Funding acquisition, Writing -original draft, Validation, Supervision, Software. Bin Qiu: Funding acquisition, Project administration, Writing-review and editing, Visualization.

## Supplementary Materials

**Supplementary Table S1**: Tibetan antelope’s data collection details.

**Supplementary Table S2**: Species annotation information for Tibetan antelope MAGs.

**Supplementary Table S3**: Antibiotic resistance gene statistics for Tibetan antelopes.

**Supplementary Table S4–S9:** Antibiotic resistance gene statistics for different species, including Tibetan ass, Tibetan cattle, Tibetan horse, Tibetan sheep, Yak, and Human, respectively.

**Supplementary Table S10:** NCBI BLAST results of shared ARGs between Tibetan antelopes and human populations.

**Supplementary Table S11:** Comprehensive information on mobile genome elements in Tibetan antelopes.

**Supplementary Table S12:** Correlation analysis between antibiotic resistance genes and mobile genetic elements in Tibetan antelopes.

**Supplementary Table S13**: Quantitative profiling of MGEs and ARGs in MAGs from Tibetan antelopes

**Supplementary Table S14**: Correlation between ARGs and VFGs

**Supplementary Table S15:** Correlation analysis between antibiotic resistance genes and virulence factors in Tibetan antelopes.

**Supplementary Table S16**: Quantitative profiling of VFGs and ARGs in MAGs from Tibetan antelopes.

**Supplementary** Fig. 1: (A) MAG processing flow. (B) Completeness and contamination of the 13,600 MAGs. (C) GC content and genome size distribution of the 13,600 MAGs. (D) Sankey diagram illustrating species annotation of these MAGs.

**Supplementary** Fig. 2: **Genomic Contribution Between VFGs and MGEs.** Circular genome maps are shown for three representative genomes. From the outermost to the innermost ring, annotations correspond to VFGs, MGEs, and ARGs, respectively. The arrow diagram illustrates the genomic co-localization patterns of ARGs, MGEs, and VFGs.

